# Intrinsically disordered protein ensembles shape evolutionary rates revealing conformational patterns

**DOI:** 10.1101/2020.07.29.227363

**Authors:** Nicolas Palopoli, Julia Marchetti, Alexander M. Monzon, Diego J. Zea, Silvio C.E. Tosatto, Maria S. Fornasari, Gustavo Parisi

## Abstract

Intrinsically disordered proteins (IDPs) lack stable tertiary structure under physiological conditions. The unique composition and complex dynamical behaviour of IDPs make them a challenge for structural biology and molecular evolution studies. Using NMR ensembles, we found that IDPs evolve under a strong site-specific evolutionary rate heterogeneity, mainly originated by different constraints derived from their inter-residue contacts. Evolutionary rate profiles correlate with the experimentally observed conformational diversity of the protein, allowing the description of different conformational patterns possibly related to their structure-function relationships. The correlation between evolutionary rates and contact information improves when structural information is taken not from any individual conformer or the whole ensemble, but from combining a limited number of conformers. Our results suggest that residue contacts in disordered regions constrain evolutionary rates to conserve the dynamic behaviour of the ensemble and that evolutionary rates can be used as a proxy for the conformational diversity of IDPs.

**Significance Statement:** Intrinsically disordered proteins (IDPs) challenge the structure-function relationship paradigm. In this work we found that individual sites of IDPs evolve under a strong rate heterogeneity, mainly due to the structural constraints imposed by contacts between their residues. This can be better explained if the contacts are taken from selected subsets of their alternative native conformations, rather than from individual conformations or the whole native ensemble. From an evolutionary point of view, this result indicates that experimentally-based ensembles are redundant. We also observed that the evolutionary rates follow the structural variability between conformers, unveiling conformational preferences. Our results set the stage for establishing novel evolutionary-based methods to study IDP ensembles.

## Introduction

The native state of proteins is composed of multiple conformers in a dynamical equilibrium, collectively known as the native ensemble^1^. Ensembles could differ in the accessibility of each conformation to the rest of conformers (limited by the interchange free energy barriers) and how much structurally different the conformers are as a whole (i.e., the conformational diversity of the protein). For globular (“ordered”) proteins or domains, the energy barriers for conformational transitions are relatively high, allowing the experimental exploration of discrete and well-established conformations with several techniques^2^. In one extreme, conformers may only account for tiny differences like those observed in the so-called rigid proteins, where slight residue movements or rotations allow the transit of ligands to binding sites, opening tunnels or enlarging cavities without backbone translations^3^. Loops movements, displacements of secondary structure elements and relative rotations of domains commonly contribute to further increase the structural differences between conformers^4^. At the other end of the protein space, in a continuous increase of conformational diversity, proteins display high plasticity and very low conformational energy barriers between conformers^5^. Consequently, these conformers are easily interchanged, difficulting the selection of a representative group that comprehensively describes the native ensemble. These proteins of large and heterogeneous conformational diversity are commonly known as Intrinsically Disordered Proteins (IDPs) or intrinsically disordered regions (IDRs). Their main characteristic is the lack of structurally well-defined conformations under native conditions^6,7^ (Figure 1A) and structurally complex native ensembles. Their distinctive properties do not prevent IDPs from being widely distributed and relatively abundant across taxa. For example, ~40% of eukaryotic proteomes^8^ and ~46% of Uniprot entries contain short disordered regions^9^. Moreover, they play a crucial role in many neurodegenerative and systemic disorders^10^.

**Figure 1.**
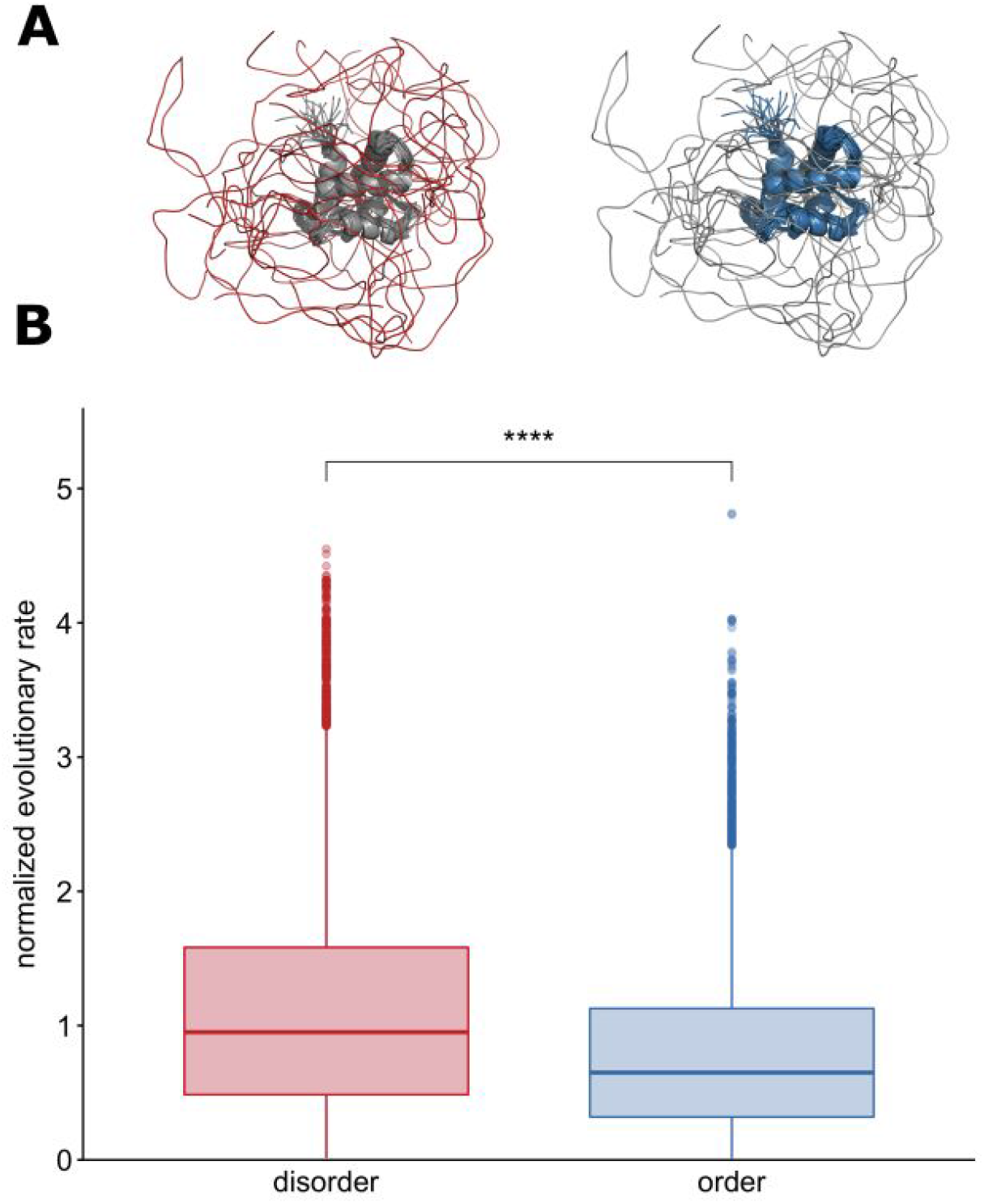
Disordered and ordered regions in intrinsically disordered proteins. **Panel A:** An example of the proteins included in our dataset. Both protein figures show the ensemble of tubulin polymerization-promoting protein family member 3 (TPPP3) from *Homo sapiens* (PDB: 2jrf), a typical IDP. Figure on the left highlights disorder regions of TPPP3 in red; such regions are included in the ‘disorder’ dataset. Figure on the right shows the TPPP3 globular domain in blue, positions that are part of the ‘order’ dataset. Proteins are represented using Pymol. **Panel B:** Boxplots showing distributions of normalized evolutionary rate per position for disordered positions of IDPs (red, median = 0.95) and for ordered positions of IDPs (blue, median = 0.65). Asterisks represent the statistical significance of the difference between a given pair of datasets (Wilcoxon rank sum test; **** indicates p <= 0.0001; ***: p<= 0.001; **: p<= 0.01; *: p <= 0.05; ns: p > 0.05)

As ordered proteins, the native structural ensemble of IDPs is a key concept in two alternative explanations describing the complex series of events associated with ligand binding^11^. In the conformational selection model, a low-populated conformation shows high binding affinity and specificity towards a given biological ligand or partner. Binding events produce a shift in the conformational equilibrium that increases the relative concentration of this competent conformation. Alternatively, the induced-fit model proposes that the adoption of the preferred conformation occurs after binding, in a process that may involve conformational changes and disorder-order transitions. Whatever the relative importance of these models to explain protein biology^12^, pre-existence of low-represented competent conformations or conformational rearrangements after ligand binding, raise the notion that IDPs ensembles are not fully disordered and that conformations with residual structure content play key biological roles^5,13–16^. One of the stronger predictors of the global evolutionary rate in proteins is the expression level of the gene^17^. However, each position evolves under different selection pressure. Several studies found correlations between per-site evolutionary rates and different structural features (for a review see^18^). For ordered proteins in general, sites with a large number of contacts, high packing densities or low solvent exposure tend to evolve more slowly^19–21^. Early studies showed that IDPs and disordered regions evolve faster than ordered proteins^22^, as expected from their low number of contacts per site^23^ and high solvent exposure^13^. Although previous works have detected structurally constrained evolutionary patterns in IDPs^24–26^, their role with IDPs typical higher flexibility remains unclear. In proteins with a significant degree of conformational diversity like IDPs, it could also be expected that different partially folded conformers with transient secondary structure elements or preferred conformations in the ensemble may impose specific and additive constraints on per-site evolutionary rates. Furthermore, these additional constraints derived from the ensemble could reveal functional types of IDPs, particularly associated with order/disorder transitions upon binding^13^. In this manuscript, we address these hypotheses by assessing the impact on evolutionary rates of the structural information derived from experimentally determined IDP ensembles.

## Results

### Evolutionary rate profiles allow the characterization of IDPs organization

Our dataset consists of data from 310 NMR ensembles of IDPs with at least 60% of disordered positions (see Methods). Figure 1A illustrates the disordered and ordered (globular) regions in these proteins. As expected, a comparison of their normalized evolutionary rate distributions (Figure 1B) shows that disordered positions of IDPs evolve significantly faster than sites from their ordered regions. This heterogeneity in evolutionary rates of IDPs is better understood by inspecting the relationship of position-specific rate profiles with their structural variation in the ensemble. We found that per-site evolutionary rates follow the structural variability between conformers measured as the RMSF (Root Mean Square Fluctuation) of their alpha carbons. Each panel in Figure 2 collects proteins that comply with a given structure-function relationship and thus have similar evolutionary rate (left column) and RMSF (middle column) profiles patterns. Cartoon depictions of representative ensembles of conformers (right column in each panel) highlight that disordered positions could occur in different regions of the protein. Disordered regions can be found in the middle or in the terminal ends of the PDB polypeptide chain (here, “terminal ends” could refer to either one of the amino or carboxyl ends of a protein or domain boundaries, as about one-third of the polypeptides in our dataset are domains occurring at least 30 residues away from the protein extremes).

**Figure 2.**
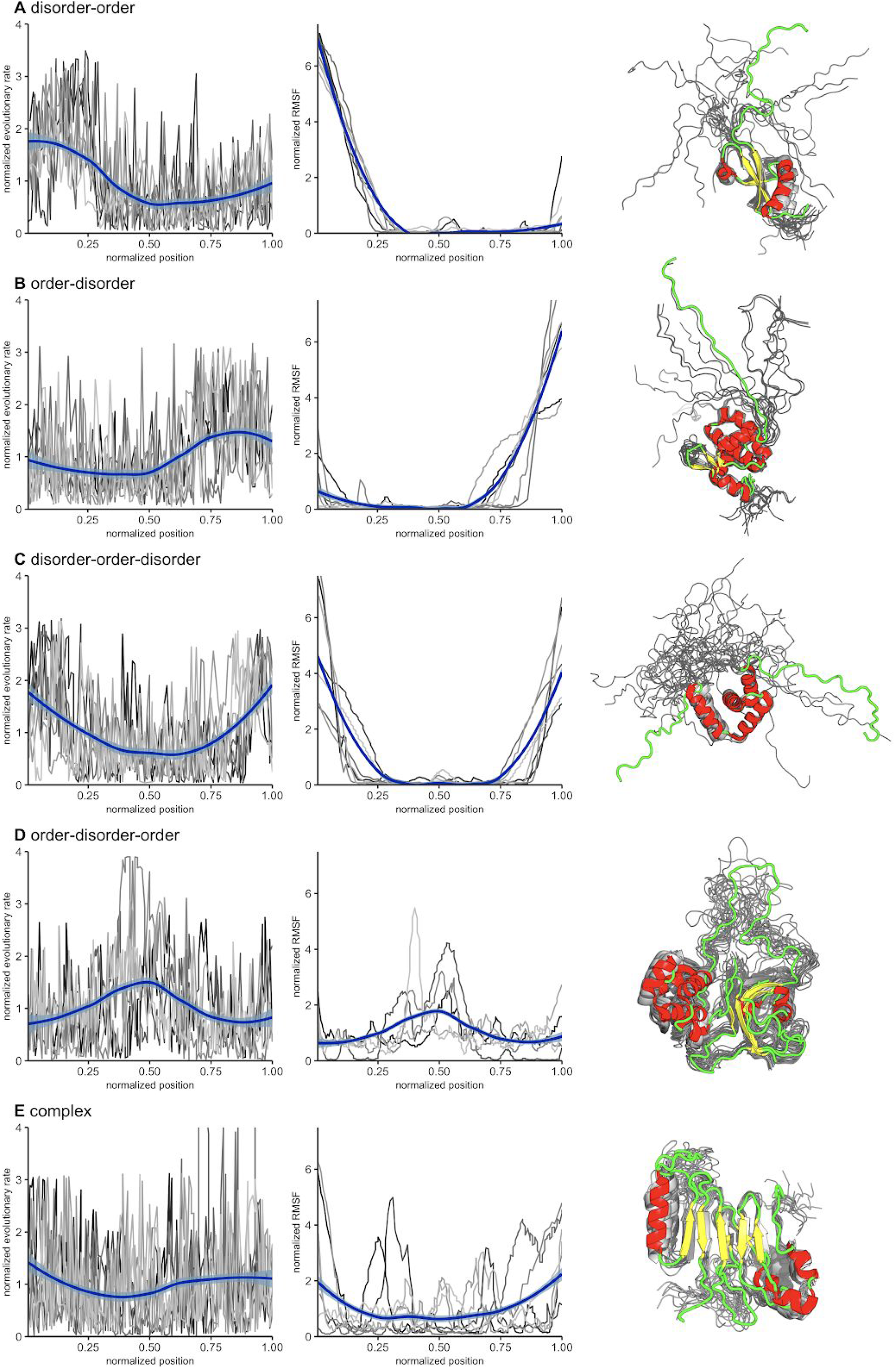
Normalized evolutionary rate profiles (left), normalized RMSF profiles (center) and cartoon image of a representative protein ensemble (right) for different structure-function patterns. Grey profiles correspond to selected protein chains from each subset. Their PDB identifiers are given below, with an asterisk marking proteins represented as cartoons (one NMR model colored by secondary structure, others in grey). Blue lines represent smooth curves obtained by LOESS regression. **Panel A**: Protein chains with a disorder-order structural pattern (PDBs: 2cqrA, 2k3aA, 2khmA, 2myxA, 2rsmA, 2rsoA, *2rvlA). **Panel B**: Proteins with an order-disorder pattern (PDBs: 1uffA, 1ug1A, 1jvrA, *2hmxA, 2k4kA, 2k5fA, 2ru8A). **Panel C**: Proteins with a disorder-order-disorder pattern (PDBs: 1ixdA, *1u6fA, 1wj7A, 2dh7A, 2ecbA, 2ejmA, 2k8pA). **Panel D**: Proteins with an order-disorder-order pattern (PDBs: 1qu6A, 2adzA, 2lbcA, 2l3sA, *2mphA). **Panel E**: Proteins with complex structural patterns (PDBs: 1mm4A, 1sm7A, 1tteA, 1vzsA, *2aivA, 2afjA, 2n1rA).

Evolutionary rate profiles can be used to distinguish at least five different protein organizations. Two of them that can be easily recognized from their rate profiles comprise an N-terminal disordered region followed by a globular domain, or vice versa (Figure 2 panels A and B, respectively). These protein chains display higher evolutionary rates in regions of high flexibility. The N-terminal domain of the murine heterochromatin protein 1 is an example of a protein with a disorder-order pattern^27^ (PDB: 2rvl; figure 2A). The N-terminal end of this domain is polyampholytic and its flexibility is regulated by phosphorylation of its serines, which changes the ensemble distribution promoting histone H3 binding. Figure 2B shows data for protein with an order-disorder organization, such as the alpha-helical MA domain from the HIV-1 Gag polyprotein precursor immediately followed by a highly flexible region (PDB: 2hmx). In the polyprotein, this disordered region links MA to the CA protein N-terminal domain. It has recently been suggested that the linker contraction may enable interactions between the immature MA and CA domains regulating the accessibility of the viral protease to release the mature proteins^28^. Some protein structures display an alternating disorder-order-disorder organization (Figure 2C). The trypanosome protein UBI-1 is one of them, with a single RNA-recognition motif domain flanked by N-terminal and C-terminal disordered regions (PDB: 1u6f)^29^. The ordered region is evidenced by a concomitant depression in both the evolutionary rate and local flexibility profiles. Other proteins have a disordered region as a linker between two globular domains (Figure 2D). As an example, the nuclear immunophilin FKBP25 (PDB: 2mph) has an N-terminal HLH domain and a C-terminal FK506-binding domain, joined by a long and flexible disordered linker that allows proper recognition of DNA by the globular domains^30^. Finally, Figure 2E combines the complex evolutionary rate and RMSF profiles derived from a heterogeneous set of protein chains. They share the presence of several short and flexible regions scattered through the sequence, disordered segments embedded in globular domains, or regions of order-disorder transitions. As an example of this structure-function relationship, the cartoon representation of the nuclear pore targeting domain of yeast nucleoporin NUP116P (PDB: 2aiv)^31^ shows regions with defined secondary structure interspersed with flexible loops.

### Effects of inter-residue contacts on evolutionary rates

Figure 1 indicates that some disordered positions can also evolve as slowly as ordered positions. To study this evolutionary rate heterogeneity in disordered regions, we estimated the average rates in disordered positions as a function of their sequence distance from the closest ordered residue in globular domains. Disordered positions have an average distance to order of^14^.4 amino acids, with a maximum of 161 amino acids. In general, disordered positions tend to evolve faster as the distance to globular regions increases. However, if bigger distances between disordered and ordered regions are considered (>20 residues), relative evolutionary rates become slower and eventually stabilize (Figure 3A). A similar behaviour is observed on the normalized values of RMSF (Figure S1). These trends might be due to the additional constraints imposed by an increased chance of establishing transient long-range interactions with distant regions in the protein, such as those responsible for multisteric regulation (for a review see^32^).

**Figure 3.**
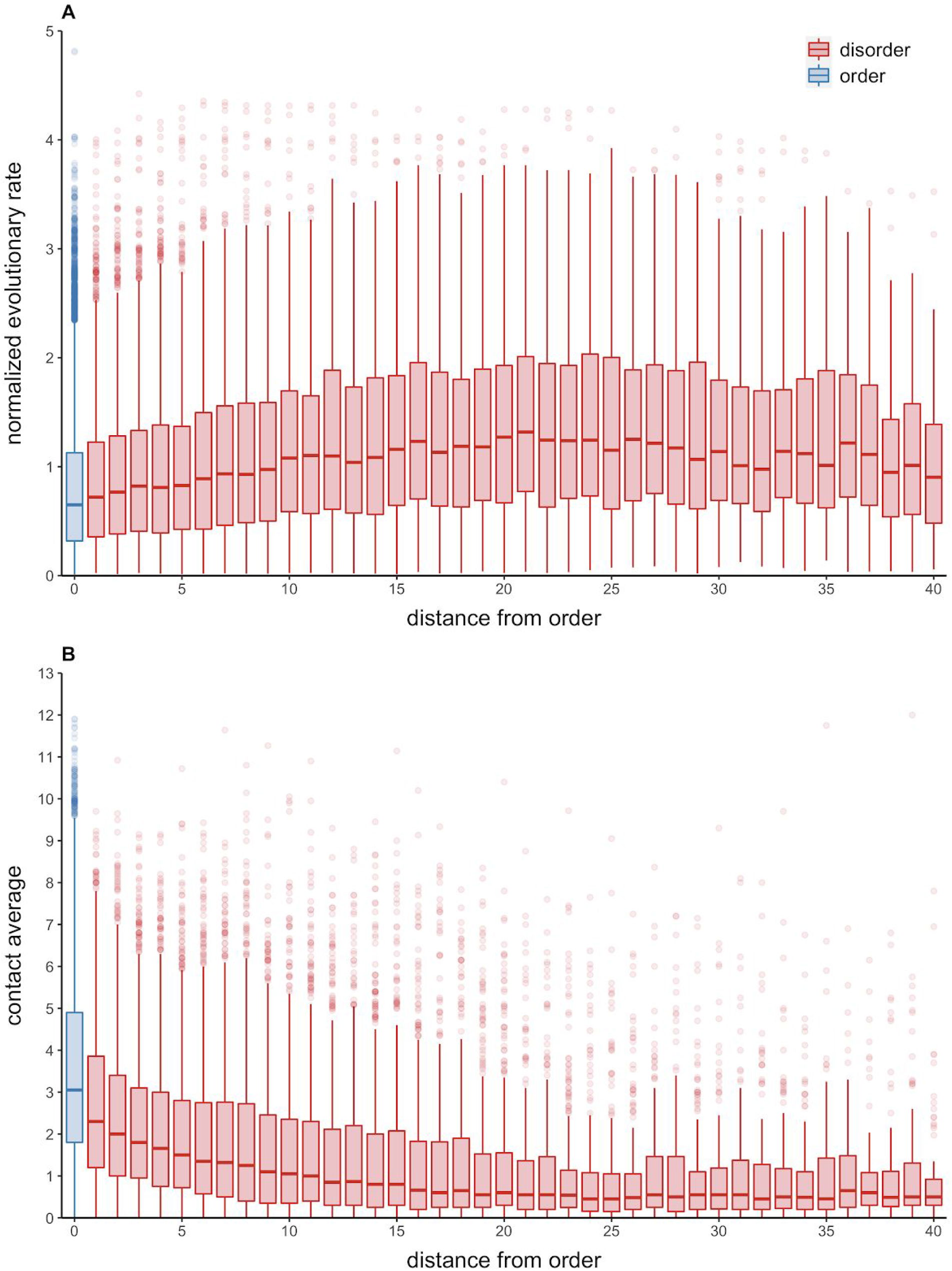
Distribution of contacts and evolutionary rates as function to distance from ordered positions. **Panel A:** Boxplots showing distribution of normalized evolutionary rates per position as function of sequence distance from the nearest ordered residue. **Panel B:** Boxplots showing distribution of average number of contacts as function of sequence distance from the nearest ordered residue.

The global evolutionary rates in Figures 1 and 2 and the rate distributions in disordered regions shown in Figure 3A are clear examples of the observed evolutionary rate heterogeneity in IDPs. The significantly different rates in ordered and disordered positions shown in Figure 1 and Figure 2 can be explained by considering the number of contacts established by each residue. The average number of contacts per position in ordered regions doubles that of disordered regions (3.44 vs 1.75, respectively). This difference evidences the higher influence of the structural constraints on the evolutionary rates in ordered regions. The influence of inter-residue contacts on rate heterogeneity extends to disordered positions. The number of contacts per site, averaged between conformers in each ensemble, decreases with their separation from ordered residues (Figure 3B). These contact number distributions show the inverse pattern to the evolutionary rates in Figure 3A.

As we mentioned above, several works studied the influence of protein structure in evolutionary rates (for a review see^18^). However, IDP ensembles are different from the globular protein ensembles analysed in those studies, since their heterogeneous and rapidly interchanging conformations make the number of residue contacts per-site to change substantially among different conformers. Considering the average of the contact number per position along all the conformers in the ensemble, we observed the same negative trend between contacts and normalized evolutionary rates regardless of the order/disorder state (Figure 4A). Interestingly, ordered and disordered regions seem to evolve at more similar rates as the number of contacts increases. We obtained similar results using other representations of per-site contacts, in particular, the maximum, minimum and mode of the number of contacts per position derived from all the conformers in the ensemble (See Supplementary Figure S2).

**Figure 4.**
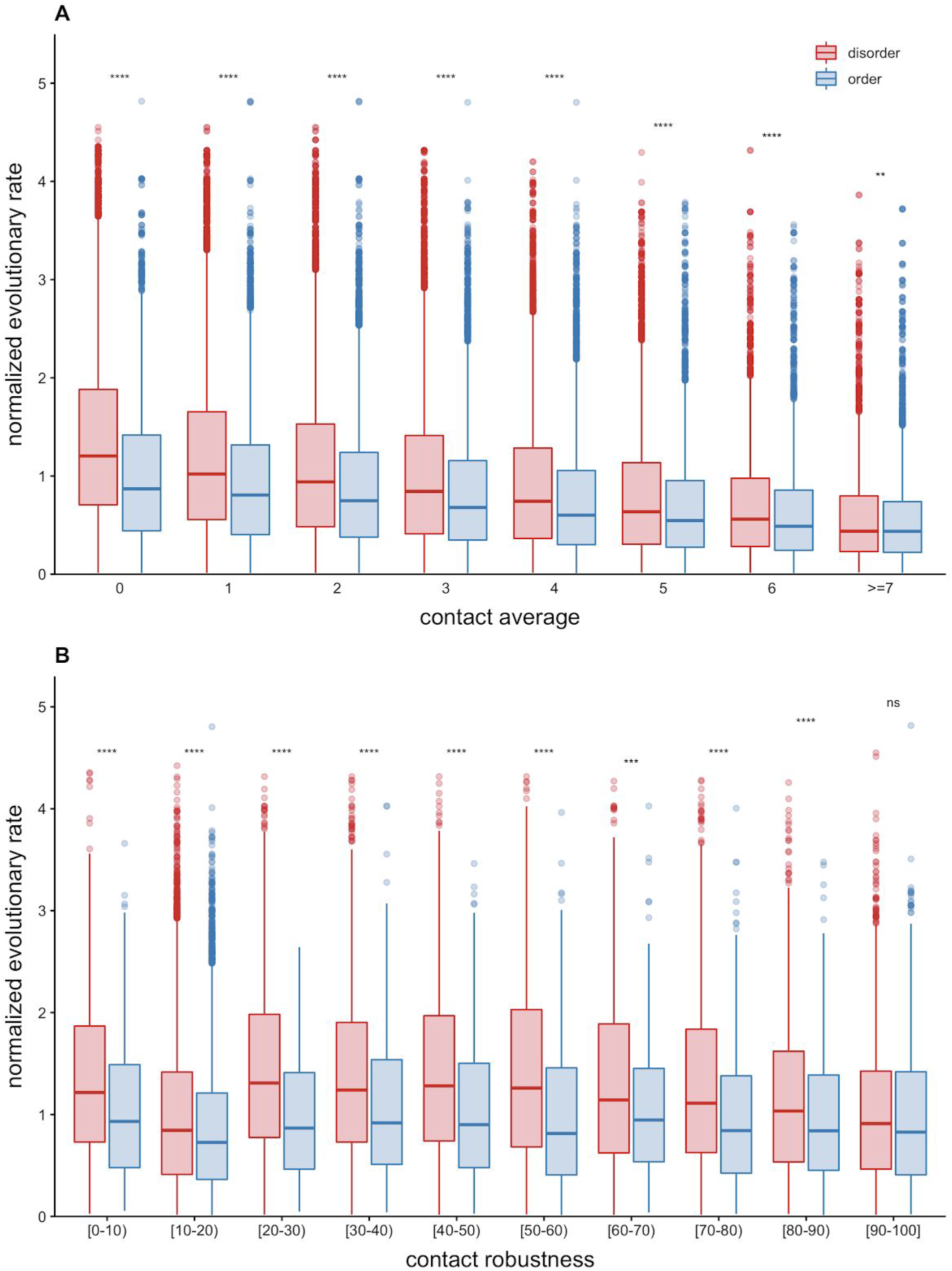
Normalized evolutionary rates variation according to contact information in the whole conformational ensemble. **Panel A:** Boxplots showing distribution of normalized evolutionary rates as function of the average number of contacts per position along the complete ensemble. **Panel B:** Boxplots showing distribution of normalized evolutionary rates as function of contact robustness. Asterisks represent the statistical significance of the difference between a given pair of datasets (Wilcoxon rank sum test; **** indicates p <= 0.0001; ***: p<= 0.001; **: p<= 0.01; *: p <= 0.05; ns: p > 0.05).

As IDPs are highly dynamical, some conformers may have a stronger structural relevance than others. We studied the relationship of evolutionary rates with the fraction of conformers having at least one contact per site in at least one conformer. For example, a value of 0.5 means that half of the observed conformers have at least one contact in that position. This fraction can be interpreted as the robustness of a position to maintain at least one contact in the ensemble; therefore, we adopted the term *contact robustness* to represent this fraction. As expected, disordered positions have lower average values of contact robustness (~0.67) than almost all the ordered positions (~0.92) (See Supplementary Figure S3). Figure 4B represents evolutionary rates as a function of contact robustness, showing that rates become slower with increasing contact robustness. Although disordered residues evolve faster regardless of the robustness, similar constraints may be acting on both disorder and ordered positions when a given residue establishes a contact in the vast majority of the conformers (contact robustness >0.9).

### Rate profiles and contacts

To further explore the relationship between conformational ensembles and evolutionary rates, we sought to determine if site-specific protein rates are better correlated with the contacts in any single conformer or with contact information derived from the whole ensemble. For each protein, the Pearson correlations of the contact average and the contact robustness per position against the evolutionary rates were calculated. Then, the best correlation for each protein among the different parameters describing the contacts was selected. For ~75% of the proteins in our dataset, the strongest correlation is observed with a single conformer from the ensemble (average ρ = −0.3692). It could be expected that conformers with the maximum number of contacts overall (assumed to be the closed form of the protein) or those with the minimum (likely the open form) may impose relevant functional constraints on evolution. However, none of these accounted for any of the best correlations in our dataset. The remaining 25% of the proteins were more associated with the contact distribution in the ensemble. Contact robustness correlations (average ρ = −0.5448) are highest for 21% of all proteins, largely surpassing the ~3% with the best correlation provided by the mean number of contacts per position (average ρ = −0.4683). Overall, for a quarter of the IDPs in our dataset, simultaneous consideration of the structural information obtained from all the available conformers in the ensemble allows for a better explanation of evolutionary change.

### Effect of conformational ensembles on evolutionary rates

In a previous work we found that IDP ensembles are redundant in terms of their structural constraints on evolution, since ~10 conformers are required on average to explain the structural constraints observed in homologous alignments of IDPs^25^. Following this idea, we explored all possible combinations of different numbers of conformers from each ensemble to study if there is a particular subset of conformers establishing contacts that better explain the evolutionary rate profiles. To this end, we first obtained all possible combinations between any number of conformers in each ensemble. Each combination was calculated as the average of residue contacts or the contact robustness per position between the selected conformers. Then, we derived the Pearson correlation coefficient rho (ρ) against the evolutionary rates by using either the contact average (ρ_ave_) or the contact robustness (ρ_crob_). We estimated the statistical significance for each combination and protein using a t-test that determines whether the obtained ρ distribution is different from the ρ distribution of a random (bootstrapped) sample (see Methods for more details). We found that in ~52% of the proteins, the evolutionary rates are better correlated with the contact robustness than with any other single conformer- or ensemble-based description of contacts (average ρ_crob_ = −0.5254). In the same sense, in ~30% of the cases, the correlations with the average of contact for a certain combination outperformed the rest (average ρ_ave_ = −0.4154). Individual conformers only provide the best correlations for ~17% of the studied proteins, and with lower ρ values (average ρ_indiv_ = −0.3695). For both the contact average and contact robustness, the correlations can be maximized by considering only a subset from the available conformers in the ensemble. The best number of combined conformers is 4.0 on average using the contact robustness and 2.8 with contact average (Figure 5).

**Figure 5.**
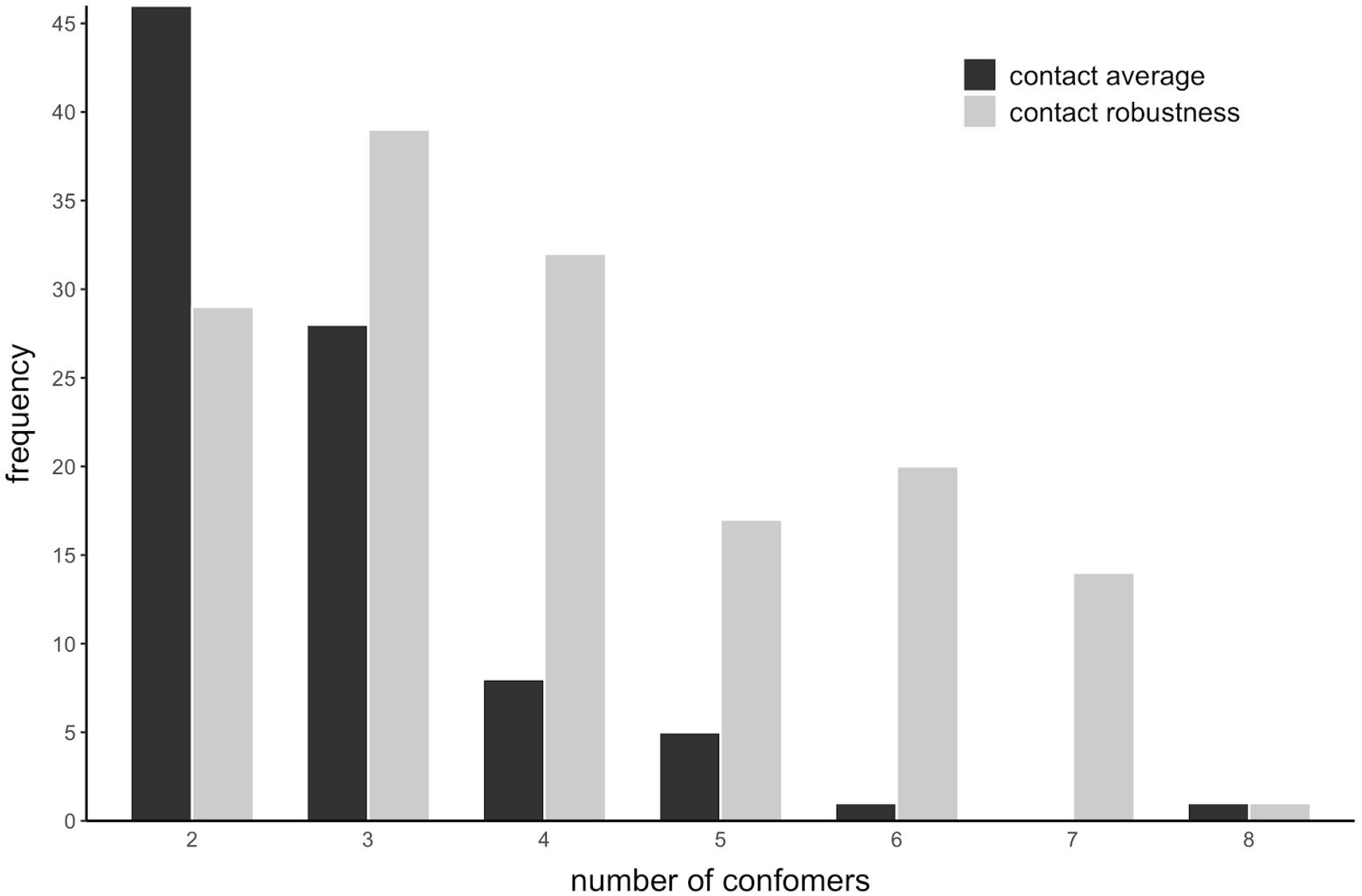
Histogram showing number of different conformers needed to find the best correlation between evolutionary rates and contact robustness (grey) or the average number of contacts (black).

The previous results highlight the importance of the ensemble in shaping evolutionary rates, as the correlations for ~82% of the proteins were improved by using different descriptions of the contacts in the conformational ensemble. Why do the evolutionary rates better correlate with the contact robustness for some proteins, while the contact average provides stronger correlations for others? While contact robustness should better represent IDPs with a considerable structural variation between conformers, contact averages could only capture the behaviour of an ensemble as a whole. We hypothesize that proteins with better ρ_crob_ (Figure 6A) could be more flexible, and their evolutionarily constrained positions could establish contacts with different positions in a given number of conformers. On the contrary, proteins better represented by ρ_ave_ (Figure 6B) might form more closed structures where the number of contacts is relatively constant along the conformers in the ensemble. To test this hypothesis, we estimated the Root Mean Square Deviation (RMSD) between all conformers in the ensemble and registered the average and the maximum. Interestingly, for the proteins for which the contact robustness provides the best correlation against evolutionary rates, the mean of the average RMSD is 19.77 Å, and the mean of their maximum RMSD is 35.2 Å. The proteins showing the best correlation against the average number of contacts per position have lower average and maximum RMSD values (with means of 11.12 Å and 20.28 Å, respectively).

**Figure 6.**
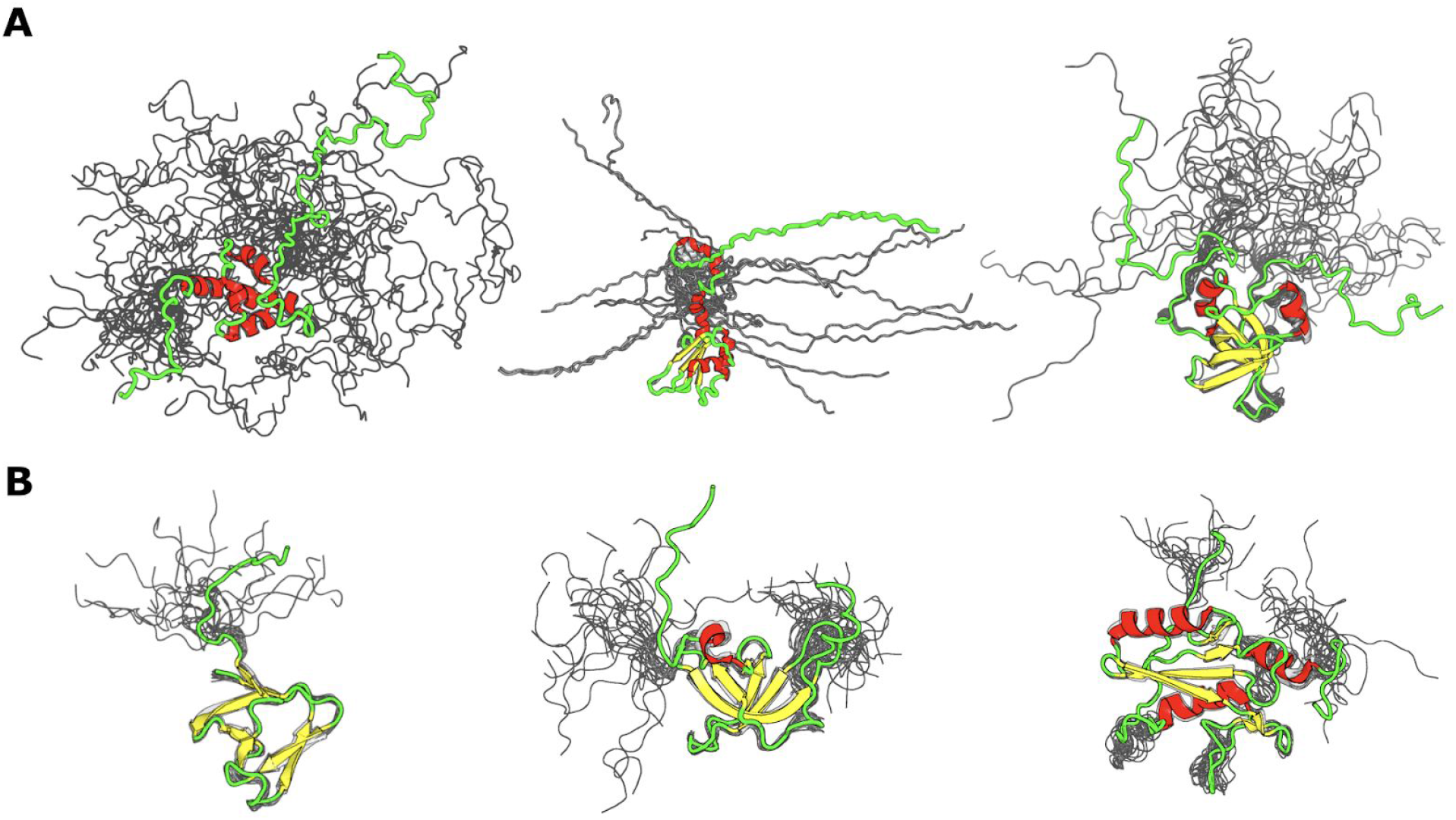
Cartoon representation of proteins for which evolutionary rates are best explained by combining contacts from selected conformers. **Panel A:** Examples of IDP ensembles for which contact robustness provides the best correlation with evolutionary rates (from left to right, PDBs: 1g9lA, 2m70A, 1d7qA). **Panel B:** Examples of IDP ensembles for which contact average correlates best with evolutionary rates (from left to right, PDBs: 2krsA, 2budA, 2eo6A).

## Discussion

In a first approximation, the evolutionary rate heterogeneity in IDPs (Figures 1, 2 and 3A) can be explained as a function of the tertiary contacts per site in different conformers of the ensemble (Figure 3B and 4). These results suggest that the state-of-the-art about evolutionary rates in globular proteins could be extended to unstructured proteins. However, IDPs share certain peculiarities. The major difference between globular proteins and IDPs is the high degree of conformational diversity in the latter and their relative ease to interchange between alternative conformers. This particular behaviour implies a marked variability in the tertiary contacts per position between the different conformers. According to our results, ~80% of the IDPs studied show better correlations between evolutionary rates and residue contact profiles when the contacts are extracted from a set of conformers instead of coming from a particular one. The best correlations are found by combining a small (<8) number of conformers rather than the whole ensemble, reflecting distinct contributions of preferred conformations, possibly related to the structure-function relationship of the protein^33,34^ These results agree with previous works describing IDP ensembles as highly redundant^25^ and open the opportunity for the development of bioinformatics tools to curate IDPs ensembles databases^35^. Largely ignored in evolutionary studies, except for a few exceptions (for example see^36–38^), contact variations due to conformational changes are revealed as impossible to neglect in IDP ensembles. These variations impose cumulative constraints on evolutionary rates^39^ that may reflect the importance of transitory or preferred conformations in the ensemble^13^.

It is possible that the observed rate heterogeneity in disordered regions may be partially influenced by the presence of functional modules involved in protein-protein interactions, ligand binding or post-translational modifications. Among these, short linear motifs (SLiMs) are typical interaction sites found in IDPs, usually more conserved than residues in their surroundings^40^. While we cannot rule these out as possible modulators of evolutionary rates in our dataset, no binding sites were found when searching for matches to known motif patterns in the ELM database^41^. Still, preferred conformations in the ensemble, detected using correlation analysis on combinations of conformers, may evidence residual secondary structures linked with their interactions.

The IDPs in our dataset have different patterns of alternating disordered and ordered regions that were evidenced by their evolutionary rate profiles. Since these profiles reflect the conformational preferences of the proteins, they could be used to infer global behaviours of IDP ensembles at the proteomic level. Interestingly, we observed the modulating effect of folded domains on the rates of attached intrinsically disordered regions, raising the question about the occurrence of coevolutionary processes^42^. Since a majority of the highest correlations with evolutionary rates were obtained using contact robustness, this supports the current view of IDPs as highly dynamic ensembles with multiple weak interactions throughout the entire protein, possibly to allow the stabilization of the ensemble and the interaction with other proteins^43–46^. In general, contact robustness reflects the evolutionary rates better than the contact number average, possibly due to the highly dynamic interactions needed to maintain the ensemble^42^.

Our results suggest that structurally-based evolutionary studies are not only feasible but also a promising approach to unveil the biology and evolution of intrinsically disordered proteins.

## Materials and Methods

### Dataset construction

We built a dataset of IDPs with known ensembles derived from Nuclear Magnetic Resonance (NMR) spectroscopy estimations deposited in the Protein Data Bank (PDB)^47^. We downloaded all PDB files from NMR experiments and selected as IDPs those that have at least 40% of their residues predicted to be disordered by Espritz^48^. We obtained pre-calculated homologous sequence alignments of these proteins from the HSSP database^49^, filtering them to allow only ~5% of gaps in average per position and with identity percentages between 33 and 100 against the PDB reference sequence (average ~60%). All remaining proteins were visually inspected, keeping only those with an evident disorder content in their 3D structure. Chimeric proteins were eliminated. This resulted in a curated dataset of 310 IDPs, with an average length of ~130 residues (minimum= 58, maximum= 370) and an average number of conformers per protein of ~20 (minimum=6, maximum=60). In this dataset, disordered regions longer than 5 residues have an average length of^23^.6 amino acids, with a maximum of 263 amino acids.

### Estimation of evolutionary rates

We fed the program Rate4Site^50^ with the alignments described above to obtain site-specific evolutionary rates for each protein. The raw (unnormalized) scores from Rate4Site were normalized for each protein separately by dividing by the protein average, following^51^. Wilcoxon rank sum tests were used to assess the statistical significance of the differences in the means of normalized evolutionary rates between ordered and disordered subsets.

### Structural information of protein conformers and ensembles

The contact information for each protein conformer was derived as the absolute number of tertiary contacts per position (defined as the minimum distances between the Van der Waals spheres of any two side-chain or *α*-carbon atoms belonging to any two amino acids, employing a cutoff distance of 1.0 Å).

Contact information for protein ensembles was estimated in several ways. Average, mode, maximum and minimum number of contacts per position were calculated over the same position in all (or a subset) of the conformers in a given ensemble. Contact robustness per position was defined as the fraction of conformers in the ensemble that establish at least one contact in that position. For example, a value of 0.5 means that half of the observed conformers have at least one contact in that position.

For each disorder position, the distance to an order position was calculated as the minimum consecutive distance to the nearest ordered residue.

Per-residue flexibility was estimated using the RSMF of the *α*-carbons in the ensemble as calculated by MIToS^52^. RMSF values were normalized by dividing them by the protein average.

### Calculation of evolutionary rate profiles and conformational patterns

For every protein in our dataset we estimated its evolutionary rate and RMSF profiles per position. As proteins have different lengths, to make proper comparisons between them, positions were renumbered to the range [0,1] by dividing by total protein length. We manually checked the order/disorder arrangement of structural elements in each protein to define its structural organization pattern. LOESS regression smoothing was performed to model the behaviour and variability of the data (normalized rates or normalized RMSF values) among selected proteins sharing the same structure-function relationship (Figure 2).

### Analysis of correlations for single conformers and for whole ensembles

For each protein conformer, we calculated the Pearson correlation coefficient *ρ* between the normalized evolutionary rates and the number of contacts per position. We repeated this on an ensemble level by deriving the Pearson correlations between rates and the contact average or contact robustness per position among all conformers of the protein. We then determined which description of contacts was better related with the evolutionary rates of each protein by comparing their correlation values.

### Analysis of correlations for combinations of subsets of conformers

For each protein and its corresponding ensemble we calculated all the possible combinations between its conformers using the formula *n*!/*k*!(*n − k*)! where *k* is the number of combined conformers and *n* is the total number of conformers for each protein. For each of these combinations, we calculated the Pearson correlation coefficient *ρ* against the evolutionary rates in two ways: by using the average of residue contacts (ρ_ave_) and by using the contact robustness (ρ_crob_). For each combination and protein, we compared each obtained ρ with its bootstrapped distribution (10,000 resamplings) to assess if each obtained ρ is significantly different from a random distribution using a t-test (significance level 0.01).

## Data and code availability

The data and code used in this manuscript are publicly available at https://gitlab.com/sbgunq/publications/palopoli-marchetti-2020-rates

## Acknowledgments

NP, MSF and GP are researchers and JM is a PhD fellow from Consejo Nacional de Investigaciones Científicas y Técnicas. AMM is funded by the research programme “Marie Skłodowska-Curie Actions Seal of Excellence @ Università degli Studi di Padova”. DJZ is funded by the Agence Nationale de la Recherche (ANR-17-CE12-0009-MASSIV). This work was supported by Universidad Nacional de Quilmes (PUNQ 1004/11), Agencia Nacional de Promoción Científica y Tecnológica (PICT-2014-3430) and the European Union’s Horizon 2020 Research and Innovation Programme (Grant Agreement N° 778247). The funders had no role in study design, data collection and analysis, decision to publish, or preparation of the manuscript.

