## Supplementary Figures for "Intrinsically disordered protein ensembles shape evolutionary rates revealing conformational patterns"

**This PDF file includes:**
Figures S1 to S3

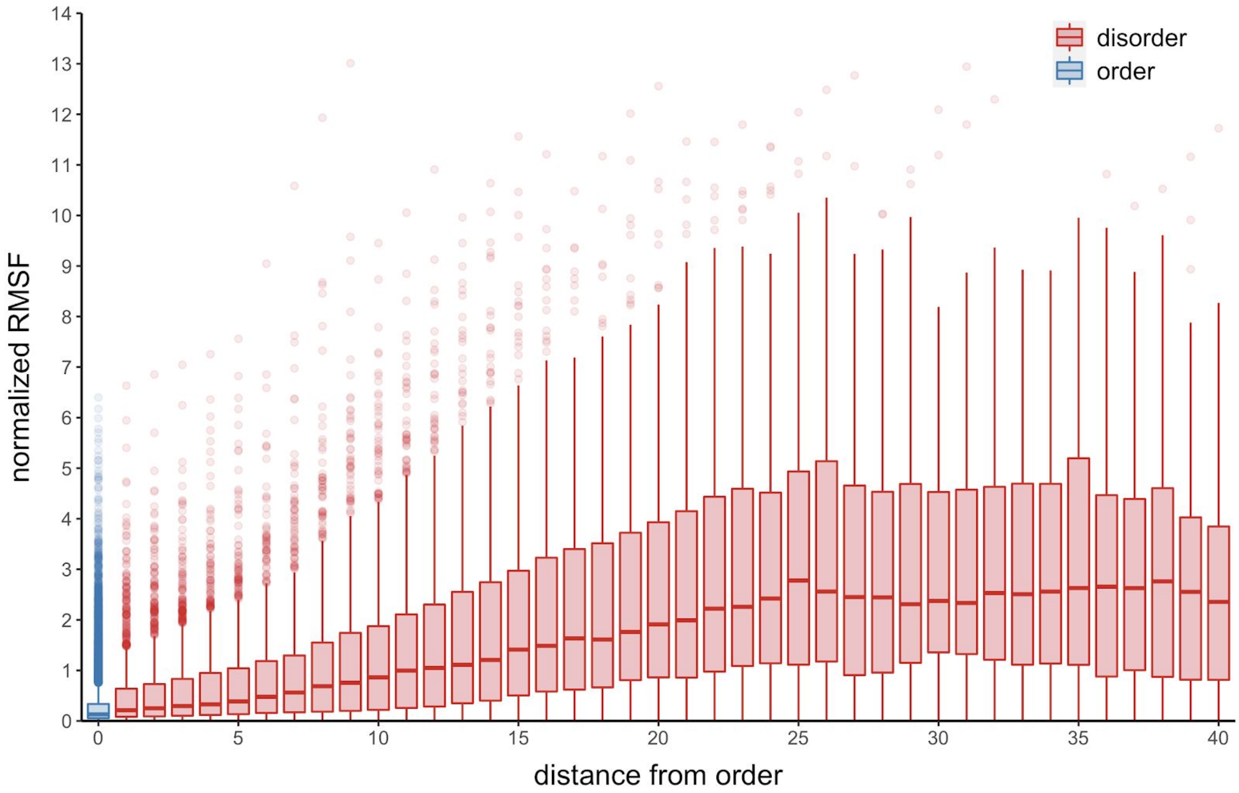


**Fig. S1.** Boxplots showing distribution of normalized RMSF values per position as function of sequence distance from the nearest ordered residue.

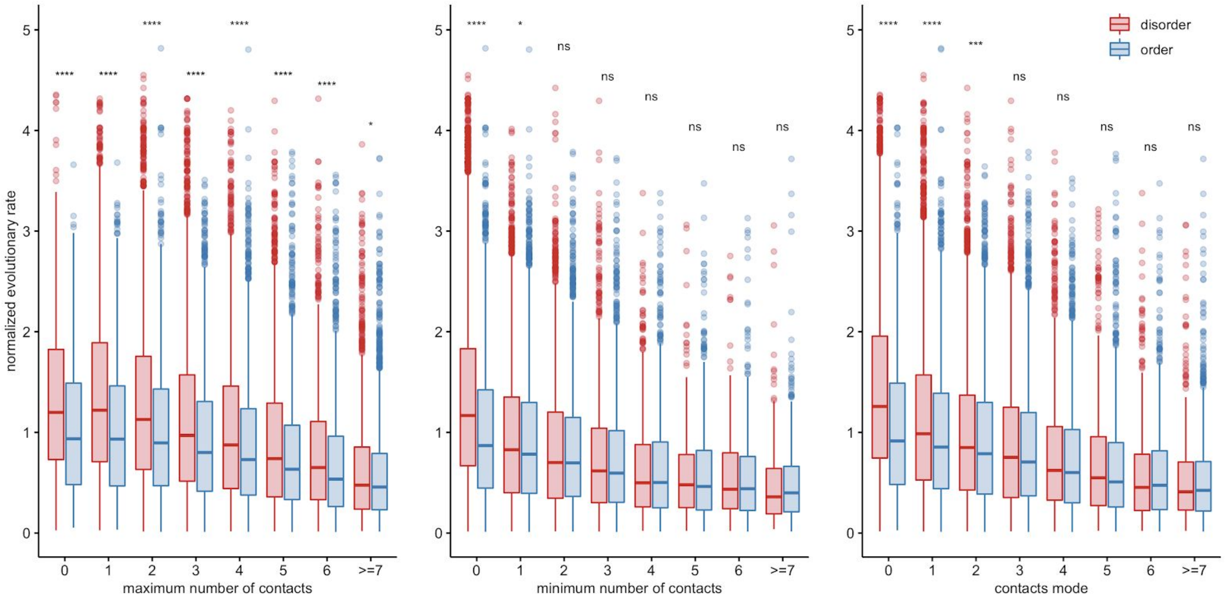


**Fig. S2.** Boxplots showing distributions of normalized evolutionary rates as function of different ensemble parameters such as maximum, minimum and mode of contacts for a given position along the ensemble.

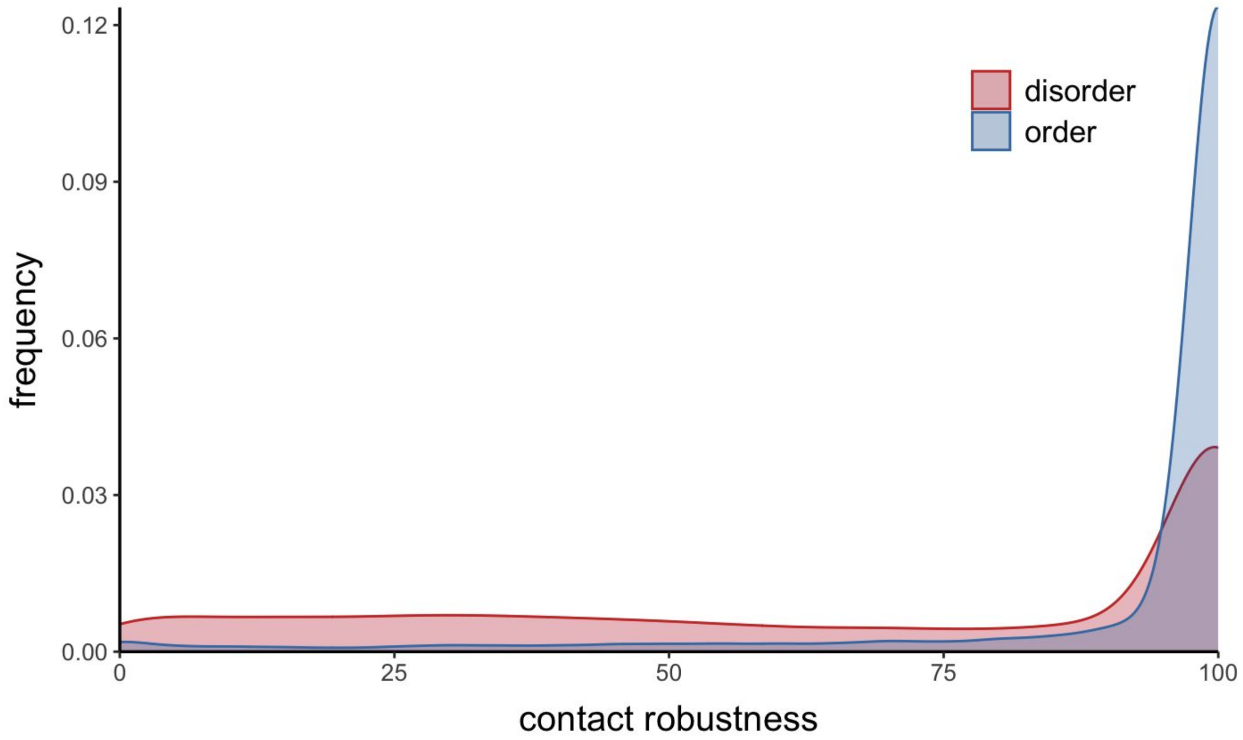


**Fig. S3.** Distribution of contact robustness for disorder (red) and order (blue) positions.
